# Roe deer embryonic diapause is associated with reduced oxidative phosphorylation and altered lipid metabolism

**DOI:** 10.64898/2026.11.15.766734

**Authors:** Sara Elsafadi, Sandra Bernal-Ulloa, Joao Agostinho de Sousa, Alexander S. Leonard, Mara D. Saenz-de-Juano, Anna-Katharina Hankele, Reto Giacometti, Thomas B. Hildebrandt, Frank Goeritz, Hubert Pausch, Ferdinand von Meyenn, Susanne E. Ulbrich

## Abstract

Conceptus elongation is a shared feature of preimplantation development in roe deer and cattle. However, unlike cattle embryos, roe deer embryos undergo embryonic diapause prior to elongation, during which blastocyst growth remains markedly reduced for 4–5 months before development rapidly resumes. To investigate the molecular basis of these developmental differences, we assembled a novel roe deer reference genome and analyzed single-embryo transcriptomes from diapausing and elongated roe deer conceptuses, comparing their transcriptomes with those of cattle embryos from day 6 morulae to day 14 tubular conceptuses. Pseudotime and principal component analyses revealed parallel developmental trajectories across species and distinct gene-expression profiles during diapause. Roe deer diapause embryos exhibited reduced transcriptional signatures of oxidative phosphorylation, mitochondrial biogenesis and protein glycosylation pathways relative to cattle blastocysts, while maintaining expression of lineage-specification markers, consistent with continued differentiation despite a metabolically quiescent state. While roe deer retained the function of the corpus luteum during diapause and continued to secrete progesterone, they did not express interferon-τ upon elongation. Elongation of roe deer embryos triggered robust upregulation of cell-cycle regulators, ribosomal biogenesis and oxidative-phosphorylation genes, aligning with the high energetic demands of elongation. By contrasting a diapause and non-diapause species, we illuminated the genomic and metabolic frameworks that underlie this unique reproductive adaptation of diapause and elongation.

## Introduction

Reproductive investment imposes a substantial metabolic burden as considerable energy is required to produce offspring (1). Approximately 130 mammalian species across diverse taxonomic orders are known to delay implantation at the blastocyst stage (2,3). This phenomenon, known as embryonic diapause, may align reproduction with favorable environmental and metabolic conditions, thereby improving offspring survival despite the substantial energetic investment made by the mother.

Embryonic diapause was first identified in the European roe deer (*Capreolus capreolus*), the only ungulate species currently known to undergo diapause (4). As seasonal breeders, roe deer breed in July–August, after which the hatched blastocysts enter a 4–5 month period of diapause, in which the blastocyst remains free-floating within the uterus proliferating at a markedly reduced rate (4). In December–January, reactivation occurs as embryonic development accelerates, and the conceptus transitions from a spherical blastocyst towards an elongated morphology before implantation. In roe deer, the postovulatory corpus luteum (CL) remains morphologically maintained. Steroidogenic activity continuous with progesterone secretion during embryonic diapause and through conceptus elongation and implantation in late December–January (5,6). Although the molecular trigger of reactivation remains incompletely defined, reactivation is associated with increased proliferation and metabolic activity, together with major changes in embryonic gene expression that become particularly apparent at elongation (4,5). The transition to elongation therefore provides a macroscopic marker of the end of diapause and subsequent resumption of rapid embryonic development (4,7).

In ruminants including roe deer and cattle, implantation is preceded by conceptus elongation, during which the hatched blastocyst transitions from an ovoid to a tubular and subsequently filamentous conceptus through extensive trophoblast proliferation and expansion (8–10). This rapid growth is accompanied by differentiation of the extraembryonic tissues and is essential for establishing sufficient maternal–conceptus contact before implantation (8–10). Conceptus elongation is highly dependent on the uterine environment and has not yet been fully recapitulated *in vitro*, although current culture systems can reproduce aspects of post-hatching development and key early events associated with elongation (11,12). This dependence is further supported *in vivo* by uterine gland knockout studies, in which loss of endometrial glands prevents normal elongation and results in early pregnancy loss (13). Cross-species embryo transfer studies further demonstrate that bovine embryos can undergo post-hatching development and elongation in a receptive ovine uterus, supporting a major role for uterine-derived signals and histotroph in regulating conceptus elongation (14). Together, these observations indicate that conceptus elongation requires coordinated maternal–conceptus interactions that provide both the regulatory cues and metabolic substrates necessary to sustain rapid trophoblast growth.

Unlike roe deer, cattle embryos develop continuously and undergo rapid conceptus elongation from approximately day 14, coinciding with increased trophoblast secretion of interferon tau (IFNτ) (15). IFNτ has long been considered the principal maternal recognition of pregnancy (MRP) signal in cattle, preventing luteolysis and allowing CL function and progesterone production to be maintained. IFNτ has also been proposed to influence conceptus elongation through autocrine and paracrine actions (16–19). However, recent evidence that bovine conceptuses lacking functional IFNτ can undergo elongation demonstrates that initiation of this developmental transition can occur independently of IFNτ (20). Consistent with this, roe deer endometrial luminal epithelial cells respond to elongated conceptuses in an IFNτ-independent manner, while CL maintenance occurs independently of pregnancy status during early gestation (21). Thus, roe deer reactivation, which marks the transition from prolonged slow development to rapid elongation, provides a natural model to identify molecular and metabolic mechanisms that act upstream of and/or independently from IFNτ. Comparative analyses with cattle may therefore reveal conserved pathways involved in initiating and sustaining successful conceptus elongation.

A key process potentially underlying developmental progression is cellular metabolism. Preimplantation mammalian embryos require substantial amounts of energy to support cell proliferation, growth and developmental progression and can utilize substrates including pyruvate, lactate, amino acids and glucose for adenosine triphosphate (ATP) generation (22). During embryonic diapause, metabolic activity is generally reduced (2), allowing the embryo to maintain viability while limiting energetic demands. Reactivation therefore requires a coordinated increase in metabolic activity to support renewed proliferation and developmental progression. In ruminants, this transition is particularly relevant because conceptus elongation is an energetically demanding process characterized by rapid trophoblast proliferation and expansion (8–10). Thus, comparing metabolic pathway regulation between diapausing and reactivated roe deer embryos and continuously developing cattle embryos may provide insight into the metabolic adaptations that enable embryonic survival during diapause and subsequent reactivation and elongation.

To enable high-resolution cross-species comparisons, we generated a *de novo* roe deer genome assembly and integrated it with existing cattle, human and cervid genomic resources to produce a highly curated *C. capreolus* annotation. Using this annotation, we performed a cross-species single-embryo transcriptomic comparison of preimplantation development in roe deer embryos collected from hunted females and cattle (*Bos taurus*) embryos produced *in vivo* across multiple developmental stages. By profiling diapausing and reactivated roe deer embryos alongside cattle embryos spanning the morula-to-elongation transition, we aimed to identify shared and species-specific molecular programs associated with diapause, reactivation and conceptus elongation.

## Methods

### Roe deer genome sequencing, assembly and annotation

We collected uterine tissue from a female roe deer and extracted genomic DNA using the Qiagen Gentra Puregene Kit. DNA was sequenced on PacBio Sequel II and IIe platforms across seven SMRT cells, totalling 114 Gb of HiFi reads with a mean read length of 19.8 Kb (NCBI: ERR11182630). We used Hifiasm to assemble a primary genome (NCBI: GCA_951849835.1 “ETH_CapCap1”), with a genome size of 3.2 Gb and 212 contigs. The contig N50 of 69.9 Mb is four orders of magnitude larger than the previous Western Roe Deer genome (NCBI: GCA_000751575.1). In the 2026-07 release build, the genome was annotated by Ensembl, with 21,343, 5,089, and 629 coding, non-coding and pseudogenes, respectively.

### Roe deer embryo collection and sequencing

Previously generated RNA-sequencing data of single roe deer embryos (Fig. 2A; Supplementary data 1) were used for this analysis (23). In brief, day 13 diapausing embryos [July] (n=6) were collected using surgical *in vivo* embryo flushing of captive roe deer (24). Additionally, diapausing [Sep–Nov] (n= 63) and elongated [Dec–Jan] (n=16) roe deer embryos were collected from regular huntings in Switzerland and Southern Germany (23). Total RNA was extracted from single roe deer embryos with the Qiagen AllPrep DNA/RNA Micro Kit (Qiagen, 80284). Due to low RNA quality and quantity, library preparation of total RNA was performed using the Clontech Pico Input Mammalian Total RNA Kit with ribosomal RNA (rRNA) depletion following the manufacturer’s instructions. Sequencing was performed by HiSeq 2500 (Illumina) at the Functional Genomics Center Zurich (FGCZ).

### *In vivo* production of cattle embryos

The animal trial was conducted in accordance with accepted standards for humane animal care and was approved by the Cantonal Veterinary Office of Zurich, Switzerland (license no. ZH042/17). The estrus cycle of adult Holstein cows was synchronized using an intravaginal controlled internal drug release dispenser (CIDR; EAZI-Breed, Zoetis, Parsippany, NJ, USA) for a period of 10 days. After the removal of the CIDR, a dose of 50 mg (2 ml) of cloprostenol (Estrumate, Merck, Animal Health, Rahway, NJ, USA) was administered intramuscularly (i.m.). A CL was confirmed through ultrasound examination after 10-11 days. To stimulate the growth of multiple follicles, hormonal treatment with porcine follicle-stimulating hormone (pFSH, Stimufol, Reprobiol SPRL, Ouffet, Belgium) began one day post CL confirmation. A total of 480 µg of FSH was injected i.m. over 4 days in 8 decreasing doses (95-95, 70-70, 45-45, and 30-30 µg) at 12-hour intervals (PM:AM). Along with the fifth FSH injection, 3 ml of Estrumate was injected. Artificial insemination (AI) took place the morning after the final FSH injection using two semen straws containing 100 million sperm (before freezing). Concurrently with AI, 5000 IU hCG (Chorulon, MSD Animal Health GmbH, Luzern, Switzerland) was administered intravenously (i.v.). Embryos were non-surgically retrieved on days 6, 8, 10, 12 and 14 (Fig. 2A; Supplementary data 1) post-AI from the uterine horns through flushing with Dulbecco’s PBS solution (Bioconcept, Allschwill, Switzerland). Subsequently, the embryos were classified based on morphology and quality according to the International Embryo Transfer Society criteria (25) then individually snap frozen for later sequencing.

### RNA-sequencing and data analysis of cattle embryos

Cattle day 6 morula (mean ± S.E.M = 165 ± 2 µm), day 8 blastocysts (181 ± 5 µm), day 10 hatched blastocysts (232 ± 29 µm), day 12 ovoid (1,382 ± 188 µm) and day 14 tubular (14,387 ± 1,818 µm) embryos were used for sequencing (*n* = 10 per group). Direct lysis was performed on day 6, day 8 and day 10 embryos. RNA extraction of day 12 and day 14 embryos was performed using Qiagen AllPrep DNA/RNA Micro Kit (Qiagen, 80284). Library preparation was performed using the Smart-seq 2 protocol coupled to poly-A selection of mRNA (26). Sequencing was performed on a Novaseq 6000 (Illumina) with 100 bp single-end reads on a full SP Flowcell at the FGCZ.

### RNA-seq alignment

Using the nf-core/rnaseq pipeline (27), RNA-seq reads from roe deer samples were aligned to the *Capreolus capreolus* reference genome assembly (GCA_013088385.1) and those from cattle were aligned to the *Bos taurus* reference genome (ARS-UCD1.2). Gene-level read counts were quantified using RSEM and imported into R using the *tximport* package (28). The quality control of the roe deer embryo libraries showed that, despite rRNA depletion prior to library preparation for sequencing, a subset of embryos showed a high rRNA percentage (> 60% rRNA) mapped (Fig. S2A). Additionally, roe deer embryos from huntings generally showed a high level of duplication (70% ->90%) (Fig. S2A). Elevated rRNA content and duplication rates were not associated with cell number, sample collection date or DNA extraction date. Therefore, prior to downstream analyses, an rRNA-related batch correction was applied to the roe deer samples. Specifically, samples were stratified into four bins according to the percentage of rRNA reads detected by *SortMeRNA* (29). These bins were treated as batch variables, and count correction was performed using *ComBat_seq* from the *sva* package (30). The cattle data was not corrected for rRNA because their levels were overall low and consistent between sample types.

### Cross-species integration

For the combined cross-species analysis, human orthologs shared between roe deer and cattle were identified, and only genes with detectable reads in both species were retained. Human gene symbols were used as a common reference. Prior to merging datasets, counts were normalized to length-scaled TPM values. An rRNA batch correction was applied again to the roe deer samples using the same four-bin strategy and *ComBat_seq* as described earlier.

### Trajectory analysis

Trajectory inference was performed on both the integrated cross-species dataset using shared human orthologs and on the roe deer samples analyzed separately. For the integrated dataset, the 500 most variable genes were selected, whereas for the roe deer–only analysis, the 2,000 most variable genes were used. Dimensionality reduction was carried out using principal component analysis (PCA), followed by *k*-means clustering. Trajectory reconstruction was then performed using *Slingshot* (31), based on the PCA embedding and cluster assignments. Each sample was assigned a pseudotime value along the inferred trajectory, which was subsequently used for trajectory visualization as well as for Gene Ontology (GO) and signaling pathway analyses.

### Differential expression and clustering

Differential gene expression analysis between developmental subcategories and species was performed using *DESeq2* (32) with a significance threshold of α = 0.05. Genes with an adjusted *p*-value < 0.05 and an absolute log₂ fold change > 2 were considered significantly differentially expressed. Expression-based clustering of these genes was performed using *Mfuzz* (33) to identify co-expression patterns.

Low-variance genes were defined separately for each species as genes that were not identified as differentially expressed between any of the analyzed developmental subcategories. Gene expression levels were then stratified into discrete abundance classes based on log₂-normalized expression values. Genes with no detectable RNA-seq counts were classified as non-expressed. Low-expression genes were defined as those with log₂-normalized counts between 0 and 3, intermediate-expression genes as those with log₂-normalized counts between 3 and 8 and high-expression genes as those with log₂-normalized counts greater than 8. This categorization was used to assess expression variability and abundance patterns independently of differential expression status.

### Dimensionality reduction and functional enrichment

PCA was performed using the 2,000 most variable genes. GO enrichment analysis was conducted using *clusterProfiler* (34) with a *p*-value cutoff of 0.05, Benjamini–Hochberg (BH) correction for multiple testing and a simplification cutoff of 0.7 to reduce redundancy among GO terms. KEGG pathway enrichment analysis was also performed using *clusterProfiler*, applying a p-value cutoff of 0.05, BH-adjusted *p*-values, nPermSimple = 10,000, and eps = 0.

### Cell cycle analysis

Cell cycle analysis was performed separately for each species. Curated lists of human S-phase and G2/M-phase marker genes were obtained from *Seurat* (35), and corresponding orthologs were identified in roe deer and cattle. For each species, gene expression data were normalized, variable features were identified, and the data were scaled prior to analysis. Cell cycle phase scores were then calculated using the *CellCycleScoring* function implemented in *Seurat*, enabling the assignment of cell cycle–related expression signatures to each sample.

### IFNτ Immunoblotting in uterine fluids

Uterine fluid (UF) from an elongated and an early implanted roe deer uterus were used to test the expression of IFNτ protein in roe deer. Cattle day 18 UF collected as described in Ulbrich et al., 2009 (36) and mammary gland cell protein extract (37) were used as positive and negative controls, respectively. Bicinchoninic acid (BCA) protein assay kit (17888228; Thermo Scientific) was used to determine protein concentration of each sample following the manufacturer’s protocol. For every sample, 20 µg of UF protein were mixed with 4x Laemmli buffer (1610747, Bio-Rad) and 2 µl of β-mercaptoethanol (1610710, Bio-Rad) and boiled at 95°C for 5 min then electrophoresed on Mini-PROTEAN® TGX™ Stain-Free 4−20% gels (4568095, Bio-Rad) at 200 V for 35 mins using Tris-Glycine-SDS buffer (A1415, BioChemia, ITW Reagents). Separated proteins were then electrically transferred (1.3 A, 7 min) to a PVDF membrane using the Trans Blot Turbo kit (1704156, Bio-Rad). Nonspecific binding was blocked in blocking solution of 5% skimmed milk (70166, Sigma-Aldrich) in 0.05% Tris buffered saline-Tween (TBST, 1706435, Bio-Rad) for 60 min and incubated with primary antibodies in blocking solution overnight at 4°C. Antibodies used were raised against bovine proteins IFNτ from two independent studies, Elmetwally et al. (38) and Schanzenbach et al. (39). After washing three times with TBST, the membrane was incubated with goat anti-rabbit/HRP conjugate (F01515, Santa-Cruz) at 1:10,000 dilution in TBST and detected with the Pierce ECL kit (Thermo Scientific, Waltham, MA). Gels and blots were analysed using a ChemiDoc MP Imaging System (Bio-Rad, Hercules, USA).

### In-gel protein digestion and protein identification by tandem mass spectrometry

Undiluted UF samples were used to maximize protein identification. Twelve microliters of pregnant D18 cattle UF (4144.3 µg/ml) and filamentous elongated pregnant roe deer UF (5330.6 µg/ml) were mixed with 4× Laemmli buffer and 2 µl β-mercaptoethanol and heated at 95°C for 5 min. Electrophoresis was performed using a Mini-PROTEAN® TGX™ 4–20% precast gel (4561096, Bio-Rad) at 200 V for 35 min. The gel was stained with Coomassie Blue (1610803, Bio-Rad) following the maximum staining intensity protocol. Gel bands corresponding to the approximate expected size of IFNτ were excised and analysed at the FGCZ where they were washed three times with 100 mM ammonium bicarbonate (NH4HCO3) in 50% acetonitrile, followed by a final wash with acetonitrile. For digestion, the gel pieces were covered with a trypsin solution in 10 mM Tris and 2 mM CaCl₂ (pH 8) and incubated overnight at 37°C. Supernatants were collected, remaining peptides were extracted with 0.1% trifluoroacetic acid (TFA) in 50% acetonitrile, and the combined supernatants were dried in a vacuum concentrator.

Digested samples were dissolved in 3% acetonitrile containing 0.1% formic acid, and peptide concentration was estimated using the Lunatic UV/Vis absorbance spectrometer (Unchained Labs). Peptides were separated on an M-class UPLC and analyzed on an Orbitrap mass spectrometer (MS; Thermo Fisher Scientific, Waltham, USA). MS data were processed using PEAKS Studio XPlus (Bioinformatic Solutions). For roe deer UF, peptide sequences were searched against the Odocoileinae database (taxonomy_id_9881) and interferon sequences from cattle (*Bos taurus*) and related Cervus species (*Cervus elaphus* and *Cervus hanglu yarkandensis*). Oxidation (M) was included as a variable modification, with no fixed modifications. Results were further analyzed using Scaffold v5.2.0 (Proteome Software Inc., Portland, USA), applying a protein false discovery rate (FDR) of 1%, a minimum of two unique peptides per identified protein and a peptide FDR of 0.1%.

### Data availability

The roe deer reference genome assembly and annotation are available through the NCBI Genome database (ETH_CapCap1; NCBI accession: GCA_951849835.1; https://www.ncbi.nlm.nih.gov/datasets/genome/GCA_951849835.1/). Previously generated roe deer embryo sequencing libraries were retrieved from the NCBI Gene Expression Omnibus (GEO; accession: GSE158806; https://www.ncbi.nlm.nih.gov/geo/query/acc.cgi?acc=GSE158806). Cattle embryo sequencing libraries are publicly available under accession number [to be added].

## Results

### Absence of interferon tau (IFNτ) in roe deer embryos

To investigate whether IFNτ, a long thought MRP signal in cattle, is conserved in roe deer, we first examined the genomic presence of *IFNT*. Using BLAST, we searched the roe deer reference genome (ETH_CapCap1) and three related cervid genomes for sequences homologous to the three functional bovine *IFNT* genes reported by Ali et al. (*IFN-tau-c1*, Gene ID: 528513; *IFNT2*, Gene ID: 317698; and *IFNT3*, Gene ID: 100313956). We identified highly similar sequences, with nucleotide identities ranging from 89.5% to 89.8% (Fig. S1). Furthermore, the genomic region containing these sequences exhibited conserved local synteny between cervids and cattle, with neighbouring genes, including MLLT3, FOCAD, HACD4, KLHL9, and ELAVL2, maintaining the same order across species. These findings indicate that *IFNT*-like sequences are present in the roe deer genome, despite evolutionary divergence between bovids and cervids.

We next investigated whether these sequences were transcriptionally active during roe deer embryonic development. Despite their genomic presence, IFNT transcripts were not detected at any of the developmental stages examined in roe deer embryos. In contrast, three IFNT isoforms were expressed in cattle embryos, with transcript levels progressively increasing during development (Fig. 1A). To determine whether IFNτ protein was present despite the absence of detectable transcripts, we analysed uterine fluid (UF) collected during roe deer conceptus elongation and implantation using immunoblotting. Two previously published antibodies known to detect bovine IFNτ, namely antibody A from Elmetwally et al. (38) and antibody B from Schanzenbach et al. (39), were used. Both antibodies detected bands of varying molecular weights in roe deer samples; however, none corresponded to the expected 22 kDa band observed for IFNτ in cattle (Fig. 1B). To assess the identity of these bands, the corresponding gel regions were subjected to mass spectrometry-based protein identification (Fig. 1C). No IFNτ-derived peptides were identified, indicating that the detected bands were likely the result of non-specific antibody binding (Supplementary Data 2).

**Figure 1:**
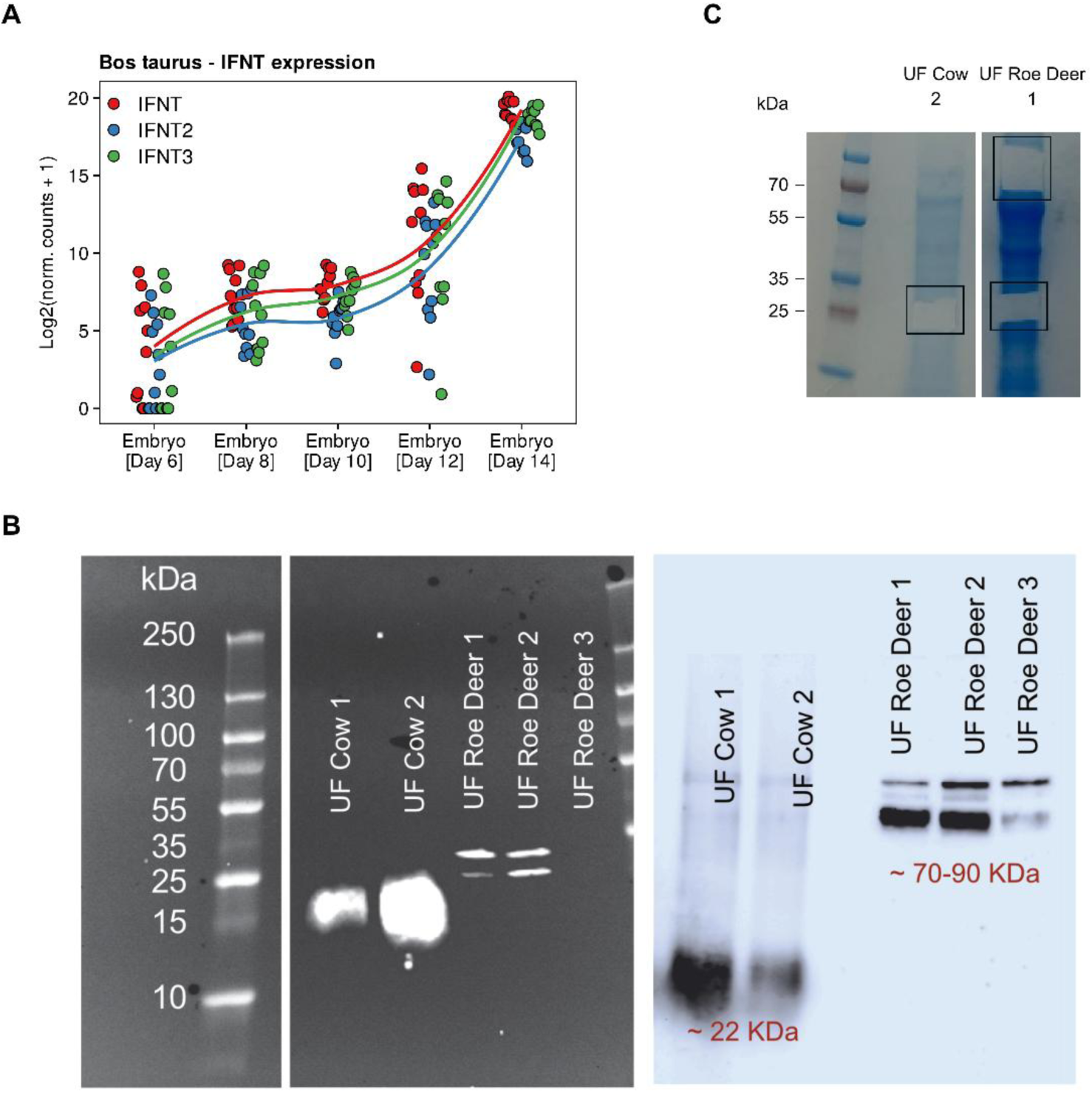
Lack of interferon tau (IFNτ) expression in roe deer peri-implantation diapause and elongated embryos. **(A)** Gene expression of three distinct IFNτ isoforms in cattle embryos. These isoforms were not expressed across the roe deer embryo stages. **(B)** Western blot of the uterine fluid of cattle elongated day 18 pregnant uterus and of elongated (UF roe deer 1 and 2) and early implanted (UF roe deer 3) roe deer pregnant uterus. Two independent anti-IFNτ antibodies were used for immunoblotting (right antibody from Elmetwally et al. (38) and left antibody from Schanzenbach et al. (39)). **(C)** Coomassie blue staining of bands detected using Antibody A and Antibody B in **(B)** used for in-gel band protein identification by mass-spectrometry.

### Inter-species transcriptomics analysis between roe deer and cattle

Given the unknown fertilisation dates of roe deer embryos collected during hunting and their slow but continuous proliferation during diapause (4), we performed a pseudotime analysis based on genes expressed across roe deer and cattle embryos (Fig. S2, S3). PCA of gene-expression profiles ordered along pseudotime revealed clear species separation along PC1 (Fig. S4A), whereas PC2 and PC3 appeared to reflect developmental progression (Fig. 2B). The expression profile of the roe deer diapausing [July] embryos closely aligned with that of the cattle day 6 morula, while the profile of the roe deer elongated [Dec–Jan] embryos closely matched that of the cattle day 14 tubular embryos (Fig. 2C). Additionally, the expression profile of the roe deer diapausing [Sep–Nov] embryos appeared to resemble that of the cattle day 10 hatched blastocysts, with notable similarities to the day 8 and day 12 cattle embryos (Fig. 2C). Projection of roe deer stage-associated gene-expression signatures onto the pseudotime PCA suggested broadly similar developmental trajectories between roe deer and cattle, while retaining species-specific expression patterns (Fig. 2D).

**Figure 2:**
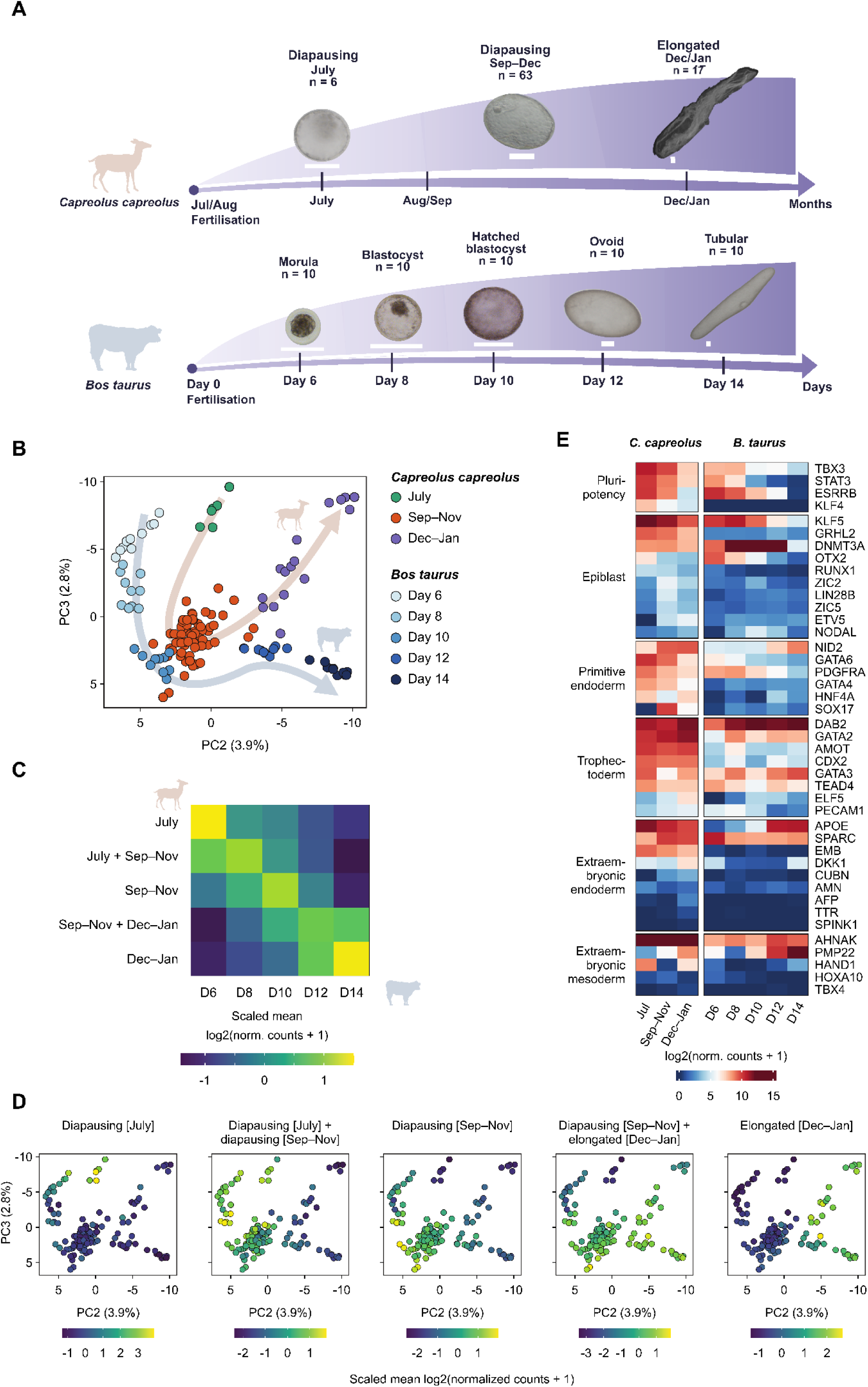
Comparative transcriptomic analysis of preimplantation embryo development in roe deer and cattle. **(A)** Illustration of roe deer and cattle preimplantation embryo development and samples used in this study. Roe deer embryos from regular huntings were grouped into diapausing [July], diapausing [Sep-Nov], and elongated [Dec-Nov] stages based on morphology and collection dates. Cattle embryos were collected on days 6, 8, 10, 12 and 14 post-fertilization via *in vivo* uterine flushing. Scale bar= 200 µm. **(B)** Pseudotime trajectory analysis of roe deer and cattle single-embryo RNA sequencing profiles. **(C)** Pseudotime analysis colour-coded by transcriptomics profiles corresponding to developmental stages in roe deer specified on top of each graph. **(D)** Correlation plot of the transcriptomics profiles between roe deer and cattle embryos. **(E)** Transcriptomics levels of key embryonic and extraembryonic markers in roe deer and cattle embryos across development stages.

We then examined the expression patterns of key embryonic and extraembryonic lineage-related genes in both species (Fig. 2E). Pluripotency-associated genes exhibited decreased expression during developmental progression in both roe deer and cattle embryos. Notably, the pluripotency markers T-box transcription factor 3 (*TBX3*) and signal transducer and activator of transcription 3 (*STAT3*) remained highly expressed in roe deer diapause embryos [Sep–Nov], whereas their expression sharply declined on day 10 in cattle embryos. Krüppel-like factor 4 (*KLF4*) showed consistently low expression in the analysed cattle embryos, whereas in roe deer its expression was higher at earlier stages and decreased with developmental progression (Fig. 2E). The epiblast marker nodal growth differentiation factor (*NODAL*) showed its highest expression in day 10 cattle embryos, while expression remained low in roe deer embryos (Fig. 2E).

Some primitive endoderm markers also exhibited distinct expression patterns between roe deer and cattle. For example, nidogen 2 (*NID2*) expression was high in both diapause [Sep–Nov] and elongated [Dec–Jan] roe deer embryos, while its expression rose only in day 12 ovoid and day 14 tubular embryos in cattle. *SRY*-box transcription factor 17 (*SOX17*) was highly expressed in roe deer diapause embryos [Sep–Nov] compared to day 8, day 10 and day 12 cattle embryos (Fig. 2E).

The trophectoderm-associated genes angiomotin (*AMOT*) and caudal type homeobox 2 (*CDX2*) seem to be highly expressed in roe deer compared to cattle embryos across the developmental stages (Fig. 2E). Interestingly, expression patterns of extraembryonic endoderm markers, including Apolipoprotein E (*APOE*), secreted protein acidic and rich in cysteine (*SPARC*) and embigin (*EMB*), differed markedly between species. *APOE* expression peaked during the earliest stage of diapause [July] and remained high across subsequent developmental stages in roe deer embryos, whereas its expression drastically increased only during conceptus elongation at days 12 and 14 in cattle embryos. *SPARC* showed an increase during both diapausing [Sep–Nov] and elongated [Dec–Jan] stages in roe deer embryos, while its expression peaked at the day 6 morula stage, subsequently declined and stabilised at lower levels till day 14 in cattle embryos. *EMB* exhibited consistently high expression in roe deer embryos but remained low across all cattle embryonic stages analysed (Fig. 2E).

### Reduced development but continued differentiation during roe deer diapause

To better understand the similarities and differences in the transcriptional landscapes of preimplantation roe deer and cattle embryos, we performed a gene set enrichment analysis on differentially expressed genes between roe deer diapause [Sep–Nov] and elongated embryos [Dec– Jan] and between day 10 hatched blastocysts and combined elongated day 12 and day 14 cattle embryos (Fig. S4B, C). We selected specific pathways to track their dynamics across roe deer pseudotime and compared them to developmental progression in cattle (Fig. 3A). Our analysis revealed that the pathways related to DNA replication, ribosome biogenesis and protein processing in the endoplasmic reticulum were downregulated in roe deer diapause [Sep–Nov] embryos as well as in cattle day 8 and day 10 embryos (Fig. 3A). Interestingly, mitochondrial transcription and translation, as well as N-glycan biosynthesis and N-linked glycosylation pathways, appeared downregulated in roe deer diapausing embryos compared to cattle blastocysts (Fig. 3A, B). In contrast, genes associated with germ cell development remained highly expressed during roe deer diapause [Sep–Nov], with a similar pattern of elevated expression in cattle day 8 and, to a lesser extent, day 10 embryos (Fig. 3A).

**Figure 3:**
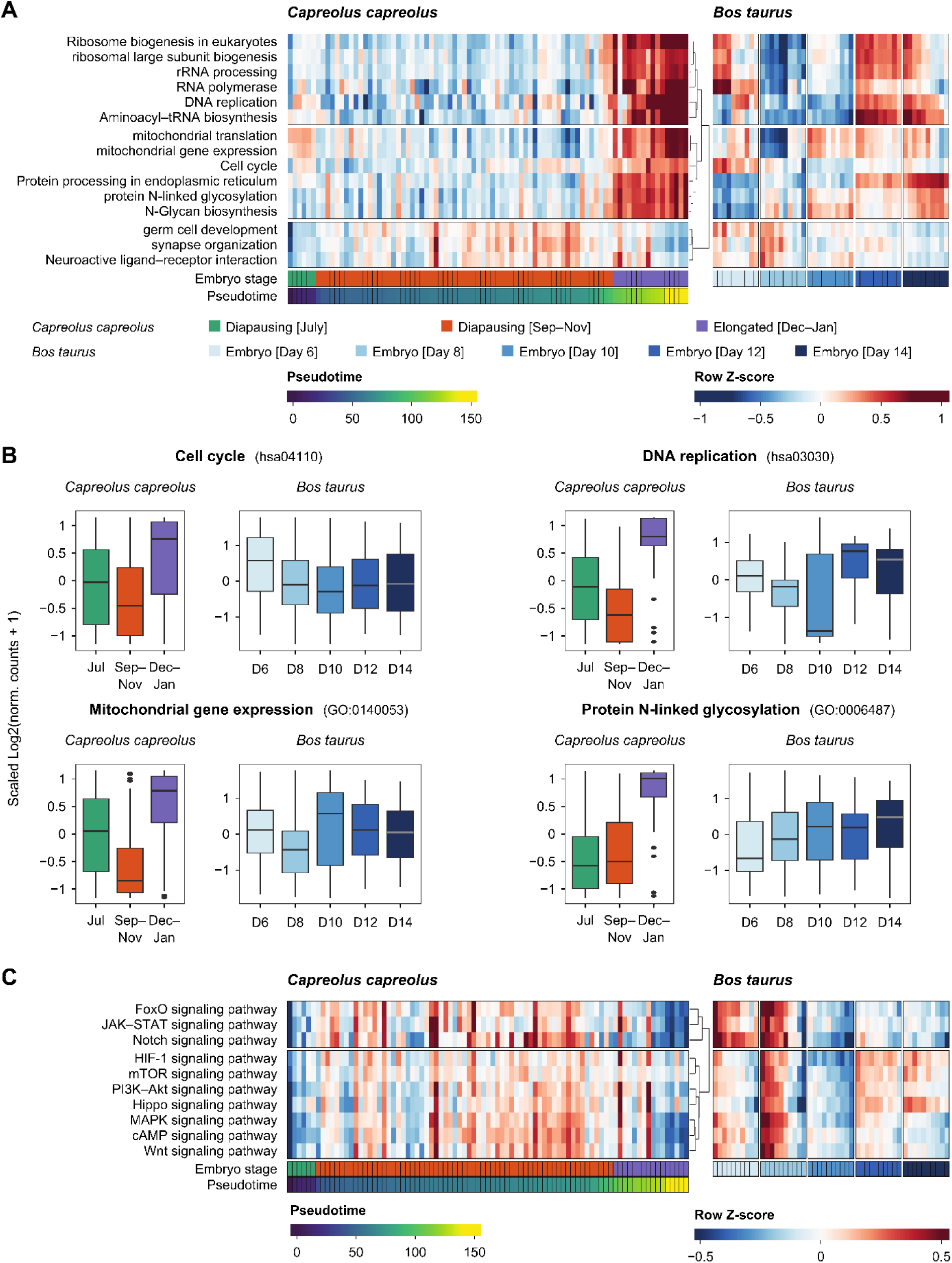
Biological processes and cell signalling status of roe deer and cattle preimplantation embryos. **(A)** Heatmap showing the raw z-score of specified gene ontology terms per embryo ranked based on pseudotime. **(B)** Expression of genes of the “Cell cycle”, “DNA replication”, “Mitochondrial gene expression” and “Protein N-linked glycosylation” gene ontology terms. **(C)** Heatmap showing the raw z-score of selected signaling pathways per embryo ranked based on pseudotime.

We further examined key signalling pathways implicated in proliferation and differentiation. In roe deer, signalling pathways including Notch, HIF-1, mTOR, Hippo, JAK-STAT, FOXO, MAPK, cAMP, and Wnt were enriched during diapause in hatched diapausing [Sep–Nov] roe deer embryos (Fig. 3C). In cattle, enrichment of these pathways peaked at the day 8 blastocyst stage but decreased in day 10 hatched blastocysts. Notably, among these pathways, HIF-1, mTOR, and Hippo signalling showed renewed enrichment in day 12 ovoid and day 14 tubular cattle embryos, corresponding to the onset of elongation, whereas in roe deer, their enrichment declined more moderately in elongated [Dec–Jan] embryos (Fig. 3C).

### Metabolism of roe deer and cattle preimplantation embryos

To examine the metabolic state of these embryos, pathway enrichment analysis indicated low involvement of oxidative phosphorylation (OxPhos) in diapause embryos [Sep–Nov], which increased in elongated embryos [Dec–Jan]. The opposite pattern was observed in cattle, where OxPhos was highly enriched in day 10 hatched blastocysts and showed lower enrichment at subsequent stages (Fig. 4A-B). Interestingly, glycolysis/glycogenesis was highly enriched upon elongation in both species. Pyruvate metabolism was also notably enriched in both species shortly before and during elongation (Fig. S4E). Fatty acid biosynthesis and degradation were implicated in elongation in both species as the raw z-score of genes involved in these pathways increased in roe deer elongated [Dec–Jan] as well day 12 and 14 cattle embryos (Fig. 4A-B, S3E). Fatty acid elongation and biosynthesis of unsaturated fatty acids likely increased shortly before and at elongation in roe deer and in cattle day 10, 12 and 14 embryos.

**Figure 4:**
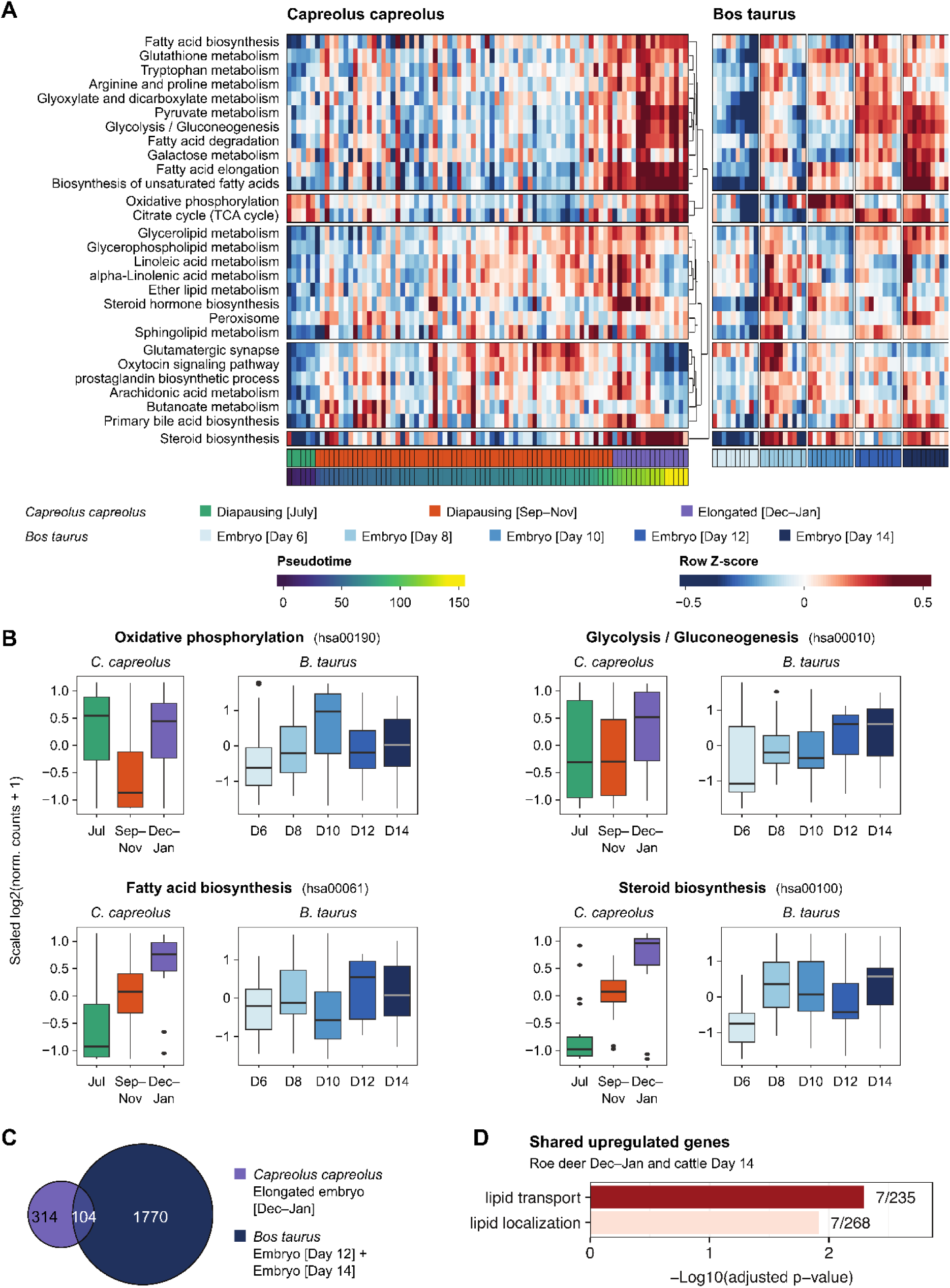
Metabolism of roe deer and cattle embryos. **(A)** Heatmap showing the raw z-scores of selected metabolic pathways involved in early embryo development and embryonic diapause. **(B)** Box plots of the expression of genes of the “Oxidative phosphorylation”, “Glycolysis/ glycogenesis”, “Fatty acid biosynthesis” and “Steroid biosynthesis” pathways in roe deer and cattle embryos. **(C)** Overlap of genes upregulated in elongated [Dc-Jan] roe deer and day 12 ovoid and 14 tubular cattle embryos. **(D)** Gene Ontology of shared genes upregulated in both species upon elongation.

Steroid biosynthesis was low during diapause but increased upon reactivation in elongated roe deer embryos. In cattle, this pathway becomes enriched at days 8 and 10 and reached its highest levels at day 14.

Differential expression analysis identified 418 upregulated genes during elongation in roe deer (diapause [Sep–Nov] vs. elongated [Dec–Jan] embryos) and 1,874 genes in cattle (day 10 vs. day 12+14 embryos). Among these, 104 genes were shared between both species (Fig. 4C; Table S1). These shared genes were significantly enriched in functions related to lipid transport and localization (Fig. 4D).

## Discussion

Our comparative transcriptomic analysis revealed that roe deer embryonic diapause is characterised by reduced transcriptional signatures of proliferation-associated and metabolic processes, including cell-cycle regulation, oxidative phosphorylation, mitochondrial biogenesis and protein post-translational glycosylation, compared with elongated embryos. Despite reduced growth, roe deer embryos maintained the expression of lineage-associated genes, consistent with continued differentiation and previously reported morphological changes (4). Cattle day 8 blastocysts similarly exhibited relatively reduced growth-and proliferation-associated transcriptional signatures while maintaining differentiation-associated signatures, potentially reflecting developmental changes associated with blastocyst hatching or differences between the stages compared. A comparative pseudotime analysis further revealed broadly similar developmental trajectories between roe deer and cattle embryos, despite their distinct reproductive strategies. However, these transcriptional similarities do not necessarily indicate equivalent developmental states, particularly given the prolonged duration of roe deer diapause. Moreover, the unknown fertilisation dates of roe deer embryos collected during hunting limit the precise determination of developmental age and the timing of molecular changes associated with diapause initiation, maintenance and exit.

Our findings support roe deer diapause as a state of reduced growth with maintained lineage integrity, further reflected by the predominance of G1-associated transcriptional signatures during later diapause and G2/M-associated signatures upon elongation (Fig. S4D). Key pluripotency factors remained expressed during roe deer diapause, while pluripotency gene expression declined with developmental progression as observed in cattle. *STAT3*, shown to be essential for inner cell mass (ICM) survival during mouse diapause (40) with anti-apoptotic activity (41), was maintained throughout roe deer diapause. Loss of *STAT3* or its receptor *Tfcp2l1* in mice has been shown to cause ICM collapse during the first few days of diapause, highlighting its conserved role (40). Similarly, *TBX3*, a gene whose homozygous deletion is embryonically lethal in mice (42) and which supports the maintenance of stem cell (43), was robustly expressed in roe deer embryos. Notably, *SOX17* was more highly expressed in roe deer diapause embryos than in days 8, 10 and 12 cattle embryos. Given its established association with the primitive endoderm/hypoblast lineage in mammalian embryos (44), this elevated expression may reflect prolonged maintenance of a primitive endoderm-associated transcriptional program during roe deer diapause. This interpretation is consistent with the previous identification of *SOX17*-positive cells and continued primitive endoderm formation in diapausing roe deer blastocysts (45). However, because these data are derived from bulk RNA-seq, it remains unclear whether the higher *SOX17* signal reflects increased expression within primitive endoderm cells, differences in lineage composition, or both. Single-cell and spatial transcriptomic approaches will therefore be required to resolve lineage-specific transcriptional programmes during diapause and reactivation.

Beyond these developmental changes, our analyses revealed pronounced species-specific differences in metabolic transcriptional programmes, particularly those associated with OxPhos. Cattle day 10 hatched blastocysts showed increased expression of OxPhos-associated genes, together with mitochondrial and pyruvate metabolism-associated signatures, although superovulation may have contributed to this pattern (46). OxPhos is active in mammalian blastocysts, including human, mouse and cattle embryos (47–49), where it supports ATP production required for blastocoel expansion (50–53). Higher OxPhos relative to glycolysis has also been associated with developmental competence (54,55), although increased mitochondrial respiration can elevate reactive oxygen species (ROS) production (54), requiring antioxidant defence mechanisms (55,56).

In contrast, roe deer diapause blastocysts exhibited reduced transcriptional signatures of OxPhos, mitochondrial transcription and mitochondrial translation, which increased again upon reactivation. These findings are consistent with a reduction in mitochondrial metabolic activity during diapause, potentially contributing to energy conservation and limiting oxidative stress (57,58), similar to the metabolic quiescence reported in dormant mouse blastocysts (58). The contrast with cattle suggests that suppression of mitochondrial metabolism may represent an adaptation to prolonged diapause rather than a general feature of the ruminant blastocyst stage. Nevertheless, prolonged diapause has also been associated with increased ROS, potentially promoting autophagic mechanisms that maintain cellular integrity and readiness for reactivation (59). As our findings are based on transcriptional signatures, direct measurements of mitochondrial respiration and ROS production will be necessary to determine the functional consequences of these changes.

The transition from diapause to rapid conceptus development is accompanied by substantial metabolic reprogramming to meet increasing energetic and biosynthetic demands. In mouse blastocysts, reactivation from diapause has been associated with enhanced glycolytic and mitochondrial activity, upregulation of pathways involved in the tricarboxylic acid (TCA) cycle and ATP synthesis and increased ATP content (58,60). A comparable increase in metabolic activity has been shown to accompany conceptus elongation in cattle, where pronounced stage-specific metabolic remodelling and increased representation of metabolites associated with glycolysis and pyruvate metabolism have been observed (61). Together, these findings suggest that enhanced glucose metabolism is a common feature of reactivation and the transition towards conceptus elongation, although specific metabolic responses may differ between species. The increased expression of glycolysis-associated genes may reflect enhanced glycolytic capacity, which could provide a rapidly accessible source of ATP while simultaneously supplying metabolic intermediates required for biosynthesis during rapid conceptus growth. Importantly, these transcriptomic patterns are consistent with our previously published UF metabolomics study, which identified stage-dependent metabolic remodelling during roe deer elongation, including changes in glycolysis and pyruvate metabolism (62). Together, these independent molecular datasets support substantial metabolic reprogramming accompanying the transition from diapause to elongation. Direct measurements of metabolic flux will be required to establish pathway activity. In addition, increased glucose utilization may support redox homeostasis through the pentose phosphate pathway, which generates NADPH required for antioxidant defence and biosynthetic processes. This may help accommodate the increased metabolic demands associated with embryonic reactivation and growth (63,64).

In addition to its role in energy production, pyruvate metabolism emerged as a key metabolism-associated pathway, linking glycolysis to mitochondrial respiration while providing carbon intermediates for fatty acid and amino acid biosynthesis (65). Our previously published UF metabolomics study further implicated the TCA cycle in citrate production for lipid synthesis, suggesting that pyruvate-related metabolism may contribute to both the energetic and lipid biosynthetic demands of conceptus elongation (62). This interpretation is further supported by our differential expression analysis, which identified 418 genes upregulated during elongation in roe deer and 1,874 in cattle, with 104 genes shared between the two species. These shared genes were significantly enriched for lipid transport and localization, indicating that remodelling of lipid handling may represent a shared feature of conceptus elongation in roe deer and cattle.

Lipid metabolism may also play an important role in maintaining embryonic viability during diapause (62,66,67). Notably, *APOE*, which encodes apolipoprotein E, a protein involved in lipid transport and metabolism (68), was highly expressed throughout roe deer diapause, with its highest expression during the earliest diapause stage analysed. In contrast, *APOE* expression increased during cattle development, reaching its highest levels during conceptus elongation at days 12 and 14. These contrasting expression patterns suggest species-specific regulation of lipid transport, potentially reflecting different metabolic requirements during prolonged diapause and continuous embryonic development. The sustained expression of *APOE* in roe deer diapause embryos may indicate an ongoing requirement for lipid handling despite reduced transcriptional signatures of mitochondrial metabolism. However, the origin of these lipids and the functional contribution of *APOE* to embryonic metabolism and survival during diapause remain to be established.

Interestingly, steroid biosynthesis followed distinct developmental trajectories between the two species. In roe deer, the expression of steroid-biosynthesis-associated genes was lower during diapause and increased following reactivation in elongated embryos, whereas in cattle, expression of steroid-biosynthesis-associated genes was already elevated at days 8 and 10 and highest at day 14. This may reflect species-specific timing of hormonal signalling and embryo–uterine interactions associated with conceptus development. Further investigation of conceptus-derived steroids, glycoproteins and uterine secretions, together with ovarian progesterone signalling, will be required to determine how these factors contribute to maintenance of pregnancy during diapause and subsequent elongation.

Beyond these metabolic adaptations, our genomic and molecular analyses revealed an additional species-specific feature of roe deer conceptus development. Analysis of our newly annotated roe deer genome identified sequences homologous to bovine IFNT genes, suggesting retention of an IFNT-related genomic region. However, no IFNT transcripts were detected at any of the roe deer embryonic stages analysed, whereas cattle embryos expressed three IFNT isoforms with increasing transcript levels during development. Conceptus-derived IFNτ has long been considered the canonical maternal recognition of pregnancy signal in ruminants (69). However, this paradigm has recently been refuted by genetic evidence from cattle, demonstrating that embryos lacking all functional IFNT copies can establish pregnancies and develop to term, resulting in the birth of healthy calves (20). Our transcriptomic, immunoblotting and proteomic findings support earlier reports of the absence of detectable IFNτ in roe deer embryos and UF. We previously demonstrated that roe deer endometrial luminal epithelial cells sense the presence of an elongated embryo in an IFNτ-independent manner (21). Together, these observations indicate that embryonic reactivation and elongation occur without detectable IFNτ production in roe deer. Roe deer herewith provide a naturally occurring ruminant model for investigating IFNτ-independent early pregnancy. It remains to be unravelled which factors mediate their maternal recognition of pregnancy, or whether, given their commonly observed monoestrous reproductive strategy, a specific embryonic signal is required at all. Similarly, the mechanisms initiating embryonic reactivation have yet to be established. Functional studies using appropriate *ex vivo* embryo–uterine models will be needed to investigate candidate metabolic pathways and alternative pregnancy signalling mechanisms.

In conclusion, our study identified distinct transcriptional profiles associated with embryonic diapause and subsequent conceptus elongation in roe deer. Its prolonged embryonic diapause was characterised by reduced OxPhos- and mitochondrial function-associated transcriptional signatures, alongside altered lipid metabolism, while lineage-associated gene expression and morphological differentiation were maintained. Reactivation and elongation were accompanied by metabolic remodelling, including increased representation of glycolytic and lipid-related pathways, some of which are shared with cattle. Given the absence of IFNτ is during early pregnancy, studying roe deer reactivation may contribute to unravelling the signal for embryo recognition and anti-luteolytic activity in other if not all ruminants. Together, these findings provide insights into the metabolic and developmental adaptations associated with embryonic diapause and establish a foundation for future functional investigations of metabolic regulation and embryo–maternal communication in ruminants.

## Abbreviations

ATP: adenosine triphosphate
AI: Artificial insemination
AMOT: Angiomotin
APOE: Apolipoprotein E
BCA: Bicinchoninic acid
CDX2: Caudal type homeobox 2
CIDR: Controlled internal drug release dispenser
CL: Corpus luteum
EMB: Embigin
FDR: False discovery rate
FGCZ: Functional Genomics Center Zurich
GO: Gene ontology
ICM: Inner cell mass
IFNτ: Interferon tau
i.m.: Intramuscularly
i.v.: Intravenously
KLF4: Krüppel-like factor 4
MS: Mass spectrometry
MRP: Maternal recognition of pregnancy
NID2: Nidogen 2
NODAL: Nodal growth differentiation factor
OxPhos: Oxidative phosphorylation
PCA: Principal component analysis
rRNA: Ribosomal RNA
ROS: Reactive oxygen species
SOX17: SRY-box transcription factor 17
SPARC: Secreted protein acidic and rich in cysteine
STAT3: Signal transducer and activator of transcription 3
TBST: Tris buffered saline-Tween
TBX3: T-box transcription factor 3
TCA: Tricarboxylic acid
TFA: Trifluoroacetic acid
UF: Uterine fluid.

## Acknowledgments

We thank the local hunters in Switzerland and Southern Germany as well as previous members of our group for their indispensable support on sample collection. We thank the Genomics and Proteomics groups at the Functional Genomics Center Zurich (FGCZ) for performing the sequencing and mass spectrometry experiments.

## References

1. Gittleman JL, Thompson SD. Energy Allocation in Mammalian Reproduction. American Zoologist. 1988;28(3): 863–875. 10.1093/icb/28.3.863.

2. Renfree MB, Fenelon JC. The enigma of embryonic diapause. Development. 2017;144(18): 3199–3210. 10.1242/DEV.148213.

3. Rüegg AB, Kowalewski MP, Ulbrich SE. Endometrial extracellular matrix components do not change over the course of embryonic diapause and reactivation in the roe deer (Capreolus capreolus). Reproduction in Domestic Animals. 2023;58(5): 594–604. 10.1111/RDA.14320.

4. Rüegg AB, Ulbrich SE. Embryonic diapause in the European roe deer – slowed, but not stopped. Animal. 2023;17: 100829. 10.1016/J.ANIMAL.2023.100829.

5. Elsafadi S, Chen H, Giacometti R, Hankele AK, Holtze S, Goeritz F, et al. Pregnanolone appearance coincides with the period of implantation following embryonic diapause in the roe deer. Reproduction. 2026;172(2): xaag091. 10.1093/reprod/xaag091.

6. Hermes R, Hildebrandt T, Goritz F, Jewgenow K, Lengwinat T, Hofmann R. Ultrasonography of the ovaries and uterus and grey scale analysis of the endometrium during embryonic diapause in European roe deer. Acta Theriologica. 2000;45(4): 559–572.

7. van der Weijden VA, Ulbrich SE. Embryonic diapause in roe deer: A model to unravel embryo-maternal communication during pre-implantation development in wildlife and livestock species. Theriogenology. 2020;158: 105–111. 10.1016/J.THERIOGENOLOGY.2020.06.042.

8. Hue I, Degrelle SA, Turenne N. Conceptus elongation in cattle: Genes, models and questions. Animal Reproduction Science. 2012;134(1–2): 19–28. 10.1016/J.ANIREPROSCI.2012.08.007.

9. Spencer TE, Forde N, Lonergan P. Insights into conceptus elongation and establishment of pregnancy in ruminants. *Reproduction*, Fertility and Development. 2017;29(1): 84–100. 10.1071/RD16359.

10. Fléchon JE, Guillomot M, Charlier M, Fléchon B, Martal J. Experimental studies on the elongation of the ewe blastocyst. Reproduction Nutrition Developpement. 1986;26(4): 1017–1024. 10.1051/RND:19860609.

11. Brandão DO, Maddox-Hyttel P, Løvendahl P, Rumpf R, Stringfellow D, Callesen H. Post Hatching Development: a Novel System for Extended in Vitro Culture of Bovine Embryos. Biology of Reproduction. 2004;71(6): 2048–2055. 10.1095/BIOLREPROD.103.025916.

12. Ramos-Ibeas P, Lamas-Toranzo I, Martínez-Moro Á, de Frutos C, Quiroga AC, Zurita E, et al. Embryonic disc formation following post-hatching bovine embryo development in vitro. Reproduction. 2020;160(4): 579–589. 10.1530/REP-20-0243.

13. Gray CA, Taylor KM, Ramsey WS, Hill JR, Bazer FW, Bartol FF, et al. Endometrial Glands Are Required for Preimplantation Conceptus Elongation and Survival1. Biology of Reproduction. 2001;64(6): 1608–1613. 10.1095/biolreprod64.6.1608.

14. Black SG, Arnaud F, Burghardt RC, Satterfield MC, Fleming JAGW, Long CR, et al. Viral Particles of Endogenous Betaretroviruses Are Released in the Sheep Uterus and Infect the Conceptus Trophectoderm in a Transspecies Embryo Transfer Model. Journal of Virology. 2010;84(18): 9078–9085. 10.1128/JVI.00950-10;PAGEGROUP:STRING:PUBLICATION.

15. Brooks K, Burns G, Spencer TE. Conceptus elongation in ruminants: Roles of progesterone, prostaglandin, interferon tau and cortisol. Journal of Animal Science and Biotechnology. 2014;5(1): 1–12. 10.1186/2049-1891-5-53.

16. Dorniak P, Bazer FW, Spencer TE. Physiology and Endocrinology Symposium: Biological role of interferon tau in endometrial function and conceptus elongation. Journal of Animal Science. 2013;91(4): 1627–1638. 10.2527/JAS.2012-5845.

17. Forde N, Lonergan P. Interferon-tau and fertility in ruminants. Reproduction. 2017;154(5): F33–F43. 10.1530/REP-17-0432.

18. Brooks K, Spencer TE. Biological roles of interferon tau (IFNT) and type I IFN receptors in elongation of the ovine conceptus. Biology of Reproduction. 2015;92(2): 1–10. 10.1095/BIOLREPROD.114.124156/2434223.

19. Bazer FW, Wu G, Johnson GA. Pregnancy recognition signals in mammals: The roles of interferons and estrogens. Animal Reproduction. 2017;14(1): 7–29. 10.21451/1984-3143-AR888.

20. Wolf E, Ali A, Zakhartchenko V, Güngör T, Nüske S, Scholz A, et al. Pregnancy establishment in cattle without embryonic interferon tau. (Pre-print). Research Square. 2026; 10.21203/rs.3.rs-8509201/v1.

21. van der Weijden VA, Puntar B, Rudolf Vegas A, Milojevic V, Schanzenbach CI, Kowalewski MP, et al. Endometrial luminal epithelial cells sense embryo elongation in the roe deer independent of interferon-tau. Biology of Reproduction. 2019;101(5): 882–892. 10.1093/BIOLRE/IOZ129.

22. Leese HJ. Metabolic control during preimplantation mammalian development. Human Reproduction Update. 1995;1(1): 63–72.

23. van der Weijden VA, Bick JT, Bauersachs S, Rüegg AB, Hildebrandt TB, Goeritz F, et al. Amino acids activate mTORC1 to release roe deer embryos from decelerated proliferation during diapause. Proceedings of the National Academy of Sciences. 2021;118(35): e2100500118. 10.1073/pnas.2100500118.

24. Drews B, Ulbrich SE, Vegas AR, Jewgenow K, Zahmel J, Roellig K, et al. Gliding into diapause: early embryo development in roe deer (Capreolus capreolus). Bioscientifica Proceedings. 2020;10: 165–180. 10.1530/biosciprocs.10.012.

25. Stringfellow DA, Givens MD, International Embryo Transfer Society. Manual of the International Embryo Transfer Society: A Procedural Guide and General Information for the Use of Embryo Transfer Technology Emphasizing Sanitary Procedures (Int Embryo Transfer Soc, Savory, IL), 4th Ed. 2010.

26. Picelli S, Faridani OR, Björklund ÅK, Winberg G, Sagasser S, Sandberg R. Full-length RNA-seq from single cells using Smart-seq2. Nature Protocols. 2014;9(1): 171–181. 10.1038/nprot.2014.006.

27. Ewels PA, Peltzer A, Fillinger S, Patel H, Alneberg J, Wilm A, et al. The nf-core framework for community-curated bioinformatics pipelines. Nature biotechnology. 2020;38(3): 276–278. 10.1038/S41587-020-0439-X.

28. Soneson C, Love MI, Robinson MD. Differential analyses for RNA-seq: Transcript-level estimates improve gene-level inferences. F1000Research. 2016;4: 1521. 10.12688/F1000RESEARCH.7563.2.

29. Kopylova E, Noé L, Touzet H. SortMeRNA: fast and accurate filtering of ribosomal RNAs in metatranscriptomic data. Bioinformatics. 2012;28(24): 3211–3217. 10.1093/BIOINFORMATICS/BTS611.

30. Zhang Y, Parmigiani G, Johnson WE. ComBat-seq: batch effect adjustment for RNA-seq count data. NAR Genomics and Bioinformatics. 2020;2(3): lqaa078. 10.1093/NARGAB/LQAA078.

31. Street K, Risso D, Fletcher RB, Das D, Ngai J, Yosef N, et al. Slingshot: cell lineage and pseudotime inference for single-cell transcriptomics. BMC Genomics. 2018;19(1): 477. 10.1186/S12864-018-4772-0.

32. Love M, Anders S, Huber W. Differential analysis of count data–the DESeq2 package. Genome Biology. 2014;15(550): 10–186. 10.1186/s13059-014-0550-8.

33. Kumar L, Futschik ME. Mfuzz: A software package for soft clustering of microarray data. Bioinformation. 2007;2(1): 5–7. 10.6026/97320630002005.

34. Yu G, Wang LG, Han Y, He QY. clusterProfiler: an R Package for Comparing Biological Themes Among Gene Clusters. Omics: a journal of integrative biology. 2012;16(5): 284–287. 10.1089/OMI.2011.0118.

35. Satija R, Farrell JA, Gennert D, Schier AF, Regev A. Spatial reconstruction of single-cell gene expression data. Nature Biotechnology. 2015;33(5): 495–502. 10.1038/nbt.3192.

36. Ulbrich SE, Schulke K, Gröbner AE, Reichenbach HD, Angioni C, Geisslinger G, et al. Quantitative characterization of prostaglandins in the uterus of early pregnant cattle. Reproduction. 2009;138(2): 371–382. 10.1530/REP-09-0081.

37. Saenz-de-Juano MD, Silvestrelli G, Bauersachs S, Ulbrich SE. Determining extracellular vesicles properties and miRNA cargo variability in bovine milk from healthy cows and cows undergoing subclinical mastitis. BMC Genomics. 2022;23(1): 189. 10.1186/S12864-022-08377-Z.

38. Elmetwally MA, Lenis Y, Tang W, Wu G, Bazer FW. Effects of catecholamines on secretion of interferon tau and expression of genes for synthesis of polyamines and apoptosis by ovine trophectoderm. Biology of Reproduction. 2018;99(3): 611–628. 10.1093/BIOLRE/IOY085.

39. Schanzenbach CI, Bernal-Ulloa SM, van der Weijden VA, Pfaffl MW, Büttner M, Wünsch A, et al. Blastocysts exhibit sex-specific signalling of IFNT transcription, translation and activity. Reproduction. 2019;157(3): 245–258. 10.1530/REP-18-0312.

40. Kraunsoe S, Azami T, Pei Y, Martello G, Jones K, Boroviak T, et al. Requirement for STAT3 and its target, TFCP2L1, in self-renewal of naïve pluripotent stem cells in vivo and in vitro. Biology Open. 2023;12(1). 10.1242/bio.059650.

41. Hirano T, Ishihara K, Hibi M. Roles of STAT3 in mediating the cell growth, differentiation and survival signals relayed through the IL-6 family of cytokine receptors. Oncogene. 2000;19(21): 2548–2556. 10.1038/sj.onc.1203551.

42. Davenport TG, Jerome-Majewska LA, Papaioannou VE. Mammary gland, limb and yolk sac defects in mice lacking Tbx3, the gene mutated in human ulnar mammary syndrome. Development. 2003;130(10): 2263– 2273. 10.1242/DEV.00431.

43. Esmailpour T, Huang T. TBX3 Promotes Human Embryonic Stem Cell Proliferation and Neuroepithelial Differentiation in a Differentiation Stage-Dependent Manner. Stem Cells. 2012;30(10): 2152–2163. 10.1002/STEM.1187.

44. Boroviak T, Loos R, Lombard P, Okahara J, Behr R, Sasaki E, et al. Lineage-Specific Profiling Delineates the Emergence and Progression of Naive Pluripotency in Mammalian Embryogenesis. Developmental Cell. 2015;35(3): 366–382. 10.1016/J.DEVCEL.2015.10.011.

45. Rüegg AB, van der Weijden VA, de Sousa JA, von Meyenn F, Pausch H, Ulbrich SE. Developmental progression continues during embryonic diapause in the roe deer. Communications Biology. 2024;7(1): 270. 10.1038/s42003-024-05944-w.

46. Gad A, Besenfelder U, Rings F, Ghanem N, Salilew-Wondim D, Hossain MM, et al. Effect of reproductive tract environment following controlled ovarian hyperstimulation treatment on embryo development and global transcriptome profile of blastocysts: implications for animal breeding and human assisted reproduction. Human Reproduction. 2011;26(7): 1693–1707. 10.1093/HUMREP/DER110.

47. Trimarchi JR, Liu L, Porterfield DM, Smith PJS, Keefe DL. Oxidative Phosphorylation-Dependent and - Independent Oxygen Consumption by Individual Preimplantation Mouse Embryos. Biology of Reproduction. 2000;62(6): 1866–1874. 10.1095/BIOLREPROD62.6.1866.

48. Lopes AS, Madsen SE, Ramsing NB, Løvendahl P, Greve T, Callesen H. Investigation of respiration of individual bovine embryos produced in vivo and in vitro and correlation with viability following transfer. Human Reproduction. 2007;22(2): 558–566. 10.1093/HUMREP/DEL404.

49. Meistermann D, Bruneau A, Loubersac S, Reignier A, Firmin J, François-Campion V, et al. Integrated pseudotime analysis of human pre-implantation embryo single-cell transcriptomes reveals the dynamics of lineage specification. Cell Stem Cell. 2021;28(9): 1625–1640.e6. 10.1016/J.STEM.2021.04.027.

50. Houghton FD, Humpherson PG, Hawkhead JA, Hall CJ, Leese HJ. Na+, K+, ATPase activity in the human and bovine preimplantation embryo. Developmental biology. 2003;263(2): 360–366. 10.1016/J.YDBIO.2003.07.014.

51. Houghton FD. Energy metabolism of the inner cell mass and trophectoderm of the mouse blastocyst. Elsevier. 2006;74(1): 11–18. 10.1111/J.1432-0436.2006.00052.X.

52. Leese HJ, Baumann CG, Brison DR, McEvoy TG, Sturmey RG. Metabolism of the viable mammalian embryo: quietness revisited. Molecular human reproduction. 2008;14(12): 667–672. 10.1093/MOLEHR/GAN065.

53. Kaneko KJ, DePamphilis ML. TEAD4 establishes the energy homeostasis essential for blastocoel formation. Development. 2013;140(17): 3680–3690. 10.1242/dev.093799.

54. Finkel T, Holbrook NJ. Oxidants, oxidative stress and the biology of ageing. Nature. 2000;408(6809): 239–247. 10.1038/35041687.

55. Betts D, Madan P. Permanent embryo arrest: molecular and cellular concepts. MHR: Basic science of reproductive medicine. 2008;14(8): 445–453. 10.1093/MOLEHR/GAN035.

56. Zhang C, Liu C, Li D, Yao N, Yuan X, Yu A, et al. Intracellular redox imbalance and extracellular amino acid metabolic abnormality contribute to arsenic-induced developmental retardation in mouse preimplantation. Journal of cellular physiology. 2010;222(2): 444–455. 10.1002/JCP.21966.

57. Fenelon JC, Banerjee A, Murphy BD. Embryonic diapause: development on hold. International Journal of Developmental Biology. 2014;58: 163–174. 10.1387/IJDB.140074BM.

58. Fu Z, Wang B, Wang S, Wu W, Wang Q, Chen Y, et al. Integral proteomic analysis of blastocysts reveals key molecular machinery governing embryonic diapause and reactivation for implantation in mice. Biology of Reproduction. 2014;90(3): 1–11. 10.1095/BIOLREPROD.113.115337/2514144.

59. Shin H, Choi S, Lim HJ. Relationship between reactive oxygen species and autophagy in dormant mouse blastocysts during delayed implantation. Clinical and Experimental Reproductive Medicine. 2014;41(3): 125–131. 10.5653/CERM.2014.41.3.125.

60. He B, Zhang H, Wang J, Liu M, Sun Y, Guo C, et al. Blastocyst activation engenders transcriptome reprogram affecting X-chromosome reactivation and inflammatory trigger of implantation. Proceedings of the National Academy of Sciences of the United States of America. 2019;116(33): 16621–16630. 10.1073/PNAS.1900401116.

61. Simintiras CA, Sánchez JM, McDonald M, O’Callaghan E, Aburima AA, Lonergan P. Conceptus metabolomic profiling reveals stage-specific phenotypes leading up to pregnancy recognition in cattle†. Biology of Reproduction. 2021;104(5): 1022–1033. 10.1093/BIOLRE/IOAB021.

62. Elsafadi S, Hankele AK, Giesbertz P, Ulbrich SE. Roe deer uterine fluid metabolome reveals elevated glycolysis, fatty acid breakdown, and spermidine synthesis upon reactivation from diapause. Biology of Reproduction. 2025;112(1): 70–85. 10.1093/BIOLRE/IOAE161.

63. Bora P, Thamodaran V, Šušor A, Bruce AW. p38-mitogen activated kinases mediate a developmental regulatory response to amino acid depletion and associated oxidative stress in mouse blastocyst embryos. bioRxiv. 2019; 807305. 10.1101/807305.

64. Kong S, Wang H. Transcriptome analysis of blastocysts acquiring implantation competency in mice. Bioscientifica Proceedings. 2020;10. 10.1530/BIOSCIPROCS.10.010.

65. Lee IK. The Role of Pyruvate Dehydrogenase Kinase in Diabetes and Obesity. Diabetes & Metabolism Journal. 2014;38(3): 181–186. 10.4093/DMJ.2014.38.3.181.

66. Ptak GE, Tacconi E, Czernik M, Toschi P, Modlinski JA, Loi P. Embryonic diapause is conserved across mammals. PLOS ONE. 2012;7(3): e33027. 10.1371/JOURNAL.PONE.0033027.

67. Arena R, Bisogno S, Gasior Ł, Rudnicka J, Bernhardt L, Haaf T, et al. Lipid droplets in mammalian eggs are utilized during embryonic diapause. Proceedings of the National Academy of Sciences of the United States of America. 2021;118(10): 2018362118. 10.1073/PNAS.2018362118.

68. Mahley RW. Apolipoprotein E: Cholesterol transport protein with expanding role in cell biology. Science. 1988;240(4852): 622–630. 10.1126/SCIENCE.3283935;WEBSITE:WEBSITE:AAAS-SITE;JOURNAL:JOURNAL:SCIENCE;WGROUP:STRING:PUBLICATION.

69. Ealy AD, Wooldridge LK. The evolution of interferon-tau. Reproduction. 2017;154(5): F1–F10. 10.1530/REP-17-0292.

