## Supplementary Figures for "Roe deer embryonic diapause is associated with reduced oxidative phosphorylation and altered lipid metabolism"

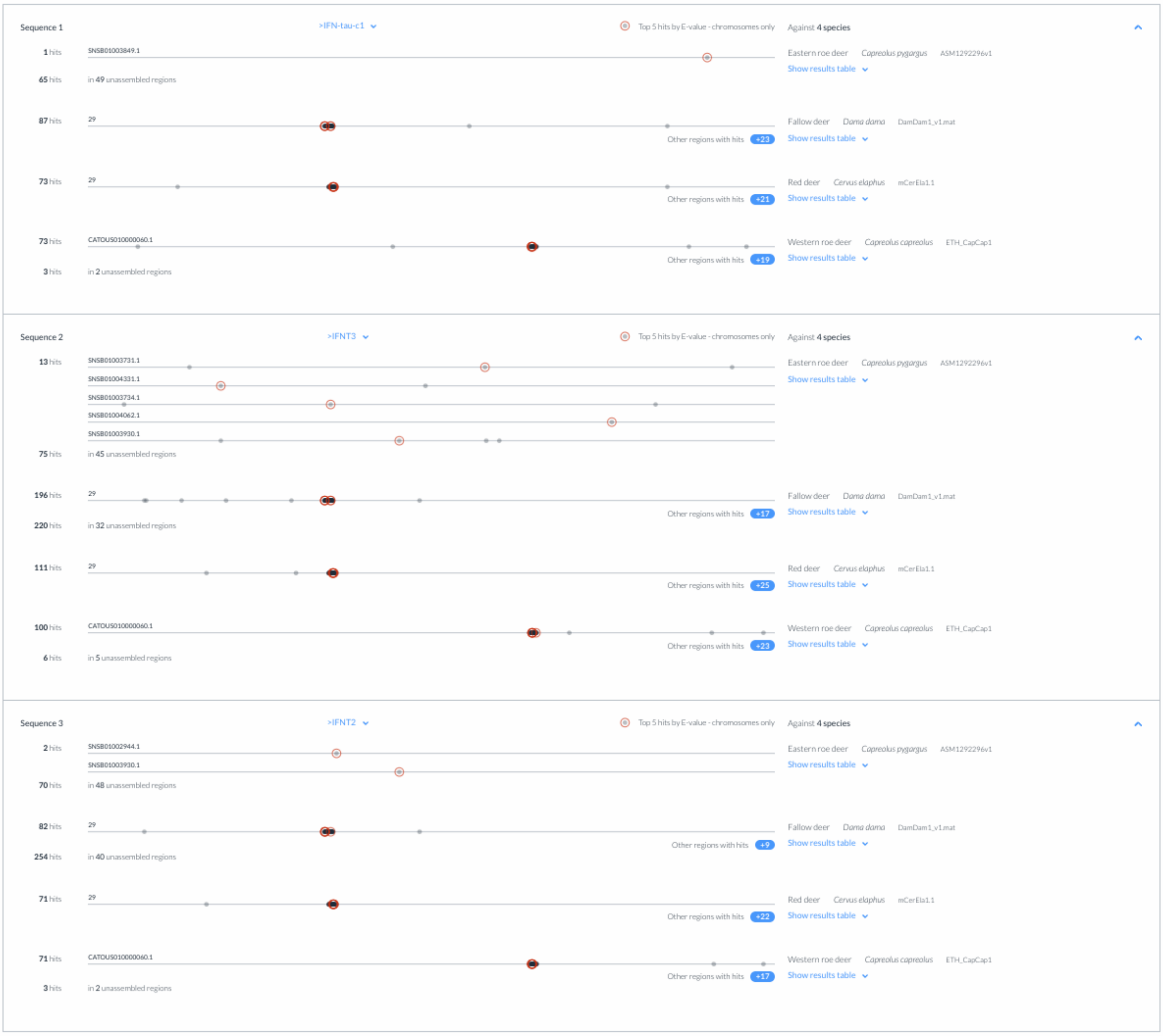


**Figure S1: Comparative genomic distribution of type I interferon genes across four cervid species.** Genomic sequence matches for *IFNA1*, *IFNT3* and *IFNT2* in Eastern roe deer, fallow deer, red deer and Western roe deer, showing the five best hits ranked by E-value.

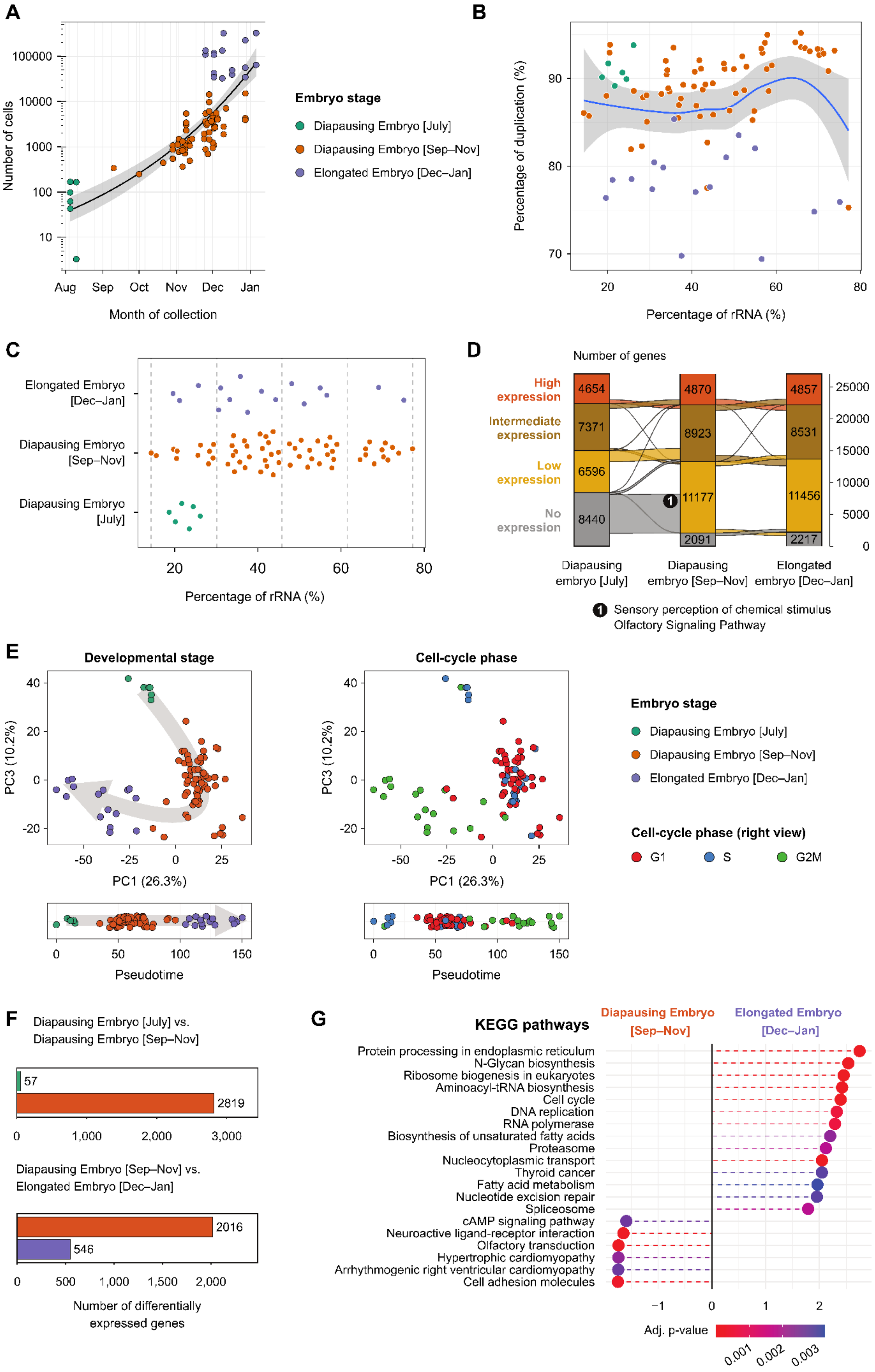
A
Number of cells
100000
10000
1000
100
10
Aug
Sep
Oct
Nov
Dec
Jan
Month of collection
Embryo stage
Diapausing Embryo [July]
Diapausing Embryo [Sep–Nov]
Elongated Embryo [Dec–Jan]
B
Percentage of duplication (%)
90
80
70
20
40
60
80
Percentage of rRNA (%)
C
Elongated Embryo [Dec–Jan]
Diapausing Embryo [Sep–Nov]
Diapausing Embryo [July]
Percentage of rRNA (%)
D
Number of genes
High expression
Intermediate expression
Low expression
No expression
4654
7371
6596
8440
4870
8923
11177
2091
4857
8531
11456
2217
25000
20000
15000
10000
5000
0
Diapausing embryo [July]
Diapausing embryo [Sep–Nov]
Elongated embryo [Dec–Jan]
1 Sensory perception of chemical stimulus Olfactory Signaling Pathway
E
Developmental stage
Cell-cycle phase
PC3 (10.2%)
PC1 (26.3%)
Pseudotime
Embryo stage
Diapausing Embryo [July]
Diapausing Embryo [Sep–Nov]
Elongated Embryo [Dec–Jan]
Cell-cycle phase (right view)
G1
S
G2M
F
Diapausing Embryo [July] vs. Diapausing Embryo [Sep–Nov]
57
2819
0
1,000
2,000
3,000
Diapausing Embryo [Sep–Nov] vs. Elongated Embryo [Dec–Jan]
2016
546
0
500
1,000
1,500
2,000
Number of differentially expressed genes
G
KEGG pathways
Diapausing Embryo [Sep–Nov]
Elongated Embryo [Dec–Jan]
Protein processing in endoplasmic reticulum
N-Glycan biosynthesis
Ribosome biogenesis in eukaryotes
Aminoacyl-tRNA biosynthesis
Cell cycle
DNA replication
RNA polymerase
Biosynthesis of unsaturated fatty acids
Proteasome
Nucleocytoplasmic transport
Thyroid cancer
Fatty acid metabolism
Nucleotide excision repair
Spliceosome
cAMP signaling pathway
Neuroactive ligand-receptor interaction
Olfactory transduction
Hypertrophic cardiomyopathy
Arrhythmogenic right ventricular cardiomyopathy
Cell adhesion molecules
−1
0
1
2
Adj. p-value
0.001
0.002
0.003

**Figure S2: Analysis of roe deer embryos sequencing data. (A)** Cell count and collection date of all roe deer samples used in this study. **(B-C)** Percentage of ribosomal RNA (rRNA) and duplication rates of roe deer sequencing data computational rRNA removal. **(D)** Transcriptomics levels across roe deer embryo stages. **(E)** Principal component analysis (PC1 and PC3) of roe deer samples in relation to embryonic stage and cell cycle. **(G)** Differentially expressed genes with gene counts reflecting those highly expressed in one stage relative to others. **(H)** Gene set enrichment analysis (GSEA) analysis of KEGG pathways comparing differential expression between diapausing (Sep–Nov) and elongated (Dec–Jan) embryos.

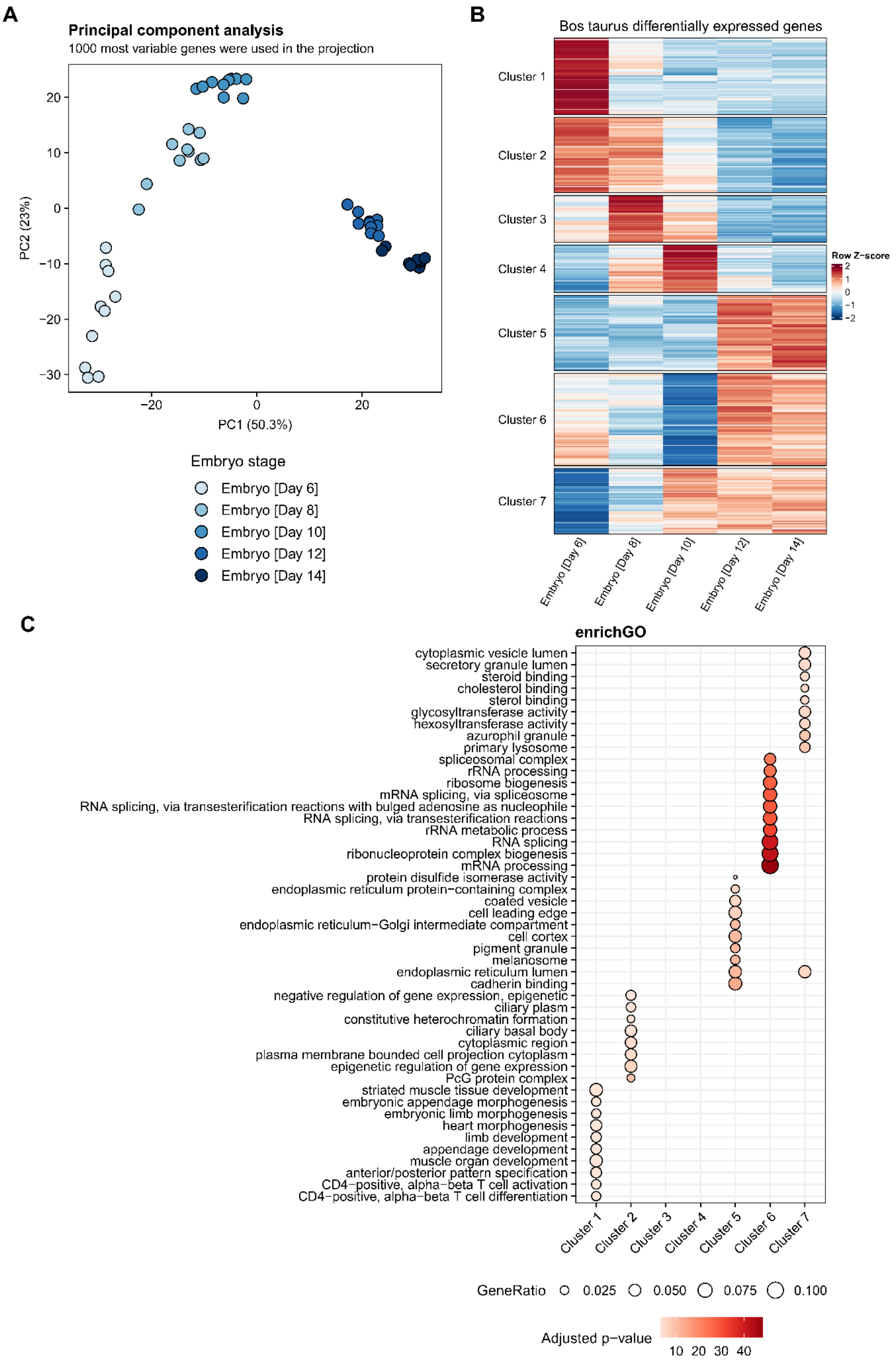
A
Principal component analysis
1000 most variable genes were used in the projection
PC2 (23%)
PC1 (50.3%)
20
10
0
−10
−20
−30
−20
0
20
Embryo stage
Embryo [Day 6]
Embryo [Day 8]
Embryo [Day 10]
Embryo [Day 12]
Embryo [Day 14]
B
Bos taurus differentially expressed genes
Cluster 1
Cluster 2
Cluster 3
Cluster 4
Cluster 5
Cluster 6
Cluster 7
Row Z−score
2
1
0
−1
−2
Embryo [Day 6]
Embryo [Day 8]
Embryo [Day 10]
Embryo [Day 12]
Embryo [Day 14]
C
enrichGO
cytoplasmic vesicle lumen
secretory granule lumen
steroid binding
cholesterol binding
sterol binding
glycosyltransferase activity
hexosyltransferase activity
azurophil granule
primary lysosome
spliceosomal complex
rRNA processing
ribosome biogenesis
mRNA splicing, via spliceosome
RNA splicing, via transesterification reactions with bulged adenosine as nucleophile
RNA splicing, via transesterification reactions
rRNA metabolic process
RNA splicing
ribonucleoprotein complex biogenesis
mRNA processing
protein disulfide isomerase activity
endoplasmic reticulum protein−containing complex
coated vesicle
cell leading edge
endoplasmic reticulum−Golgi intermediate compartment
cell cortex
pigment granule
melanosome
endoplasmic reticulum lumen
cadherin binding
negative regulation of gene expression, epigenetic
ciliary plasm
constitutive heterochromatin formation
ciliary basal body
cytoplasmic region
plasma membrane bounded cell projection cytoplasm
epigenetic regulation of gene expression
PcG protein complex
striated muscle tissue development
embryonic appendage morphogenesis
embryonic limb morphogenesis
heart morphogenesis
limb development
appendage development
muscle organ development
anterior/posterior pattern specification
CD4−positive, alpha−beta T cell activation
CD4−positive, alpha−beta T cell differentiation
Cluster 1
Cluster 2
Cluster 3
Cluster 4
Cluster 5
Cluster 6
Cluster 7
GeneRatio 0.025 0.050 0.075 0.100
Adjusted p−value
10 20 30 40

**Figure S3: Analysis of cattle embryos sequencing data. (A)** Principal component analysis (PCA) of bovine embryo transcriptomics data across developmental stages. **(B)** Mfuzz analysis across cattle embryo stages. **(C)** Gene Ontology (GO) enrichment analysis of the gene clusters identified in (B), performed using enrichGO.

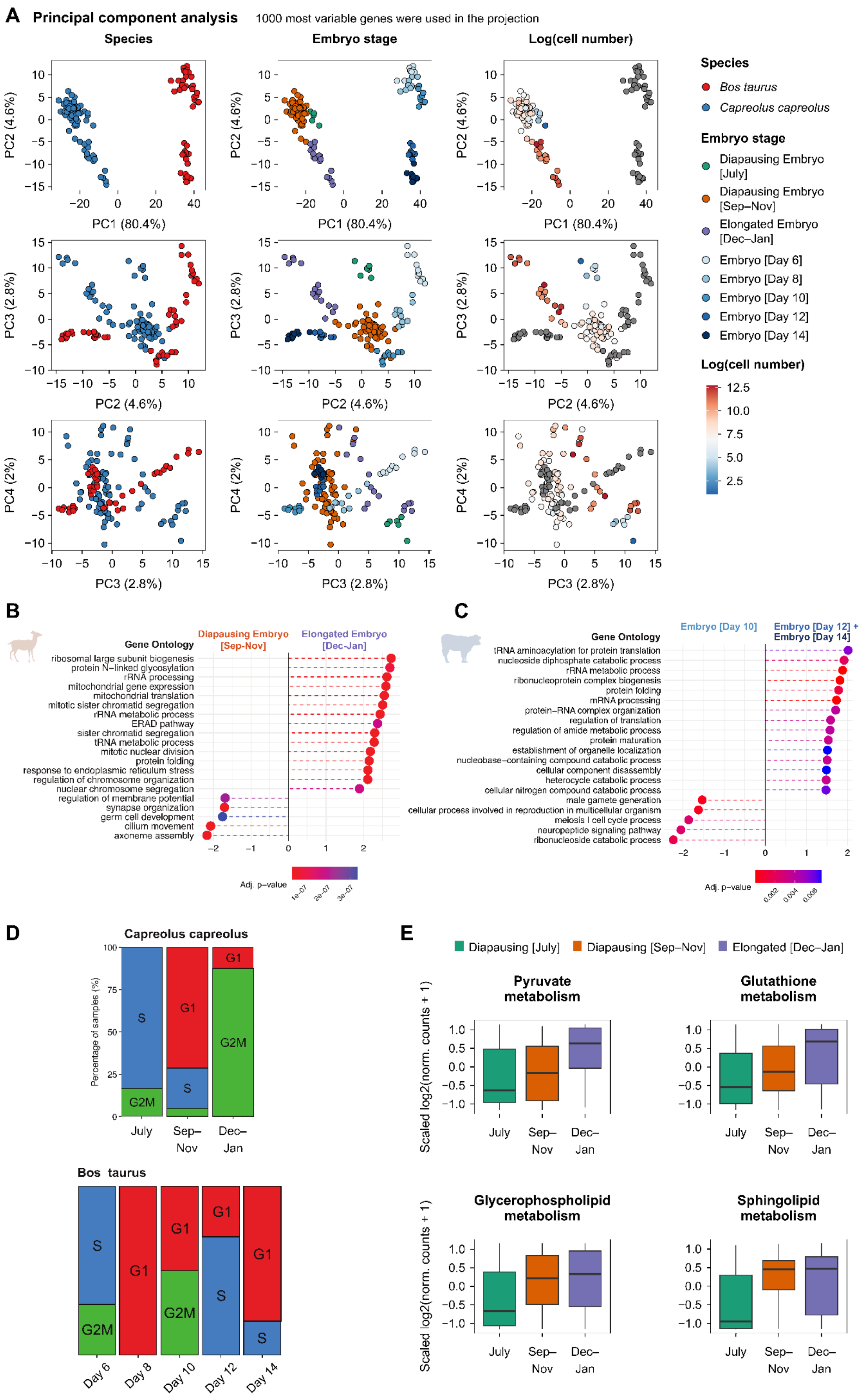

A Principal component analysis
1000 most variable genes were used in the projection
Species
Embryo stage
Log(cell number)
PC1 (80.4%)
PC2 (4.6%)
PC3 (2.8%)
PC4 (2%)
Species
Bos taurus
Capreolus capreolus
Embryo stage
Diapausing Embryo [July]
Diapausing Embryo [Sep–Nov]
Elongated Embryo [Dec–Jan]
Embryo [Day 6]
Embryo [Day 8]
Embryo [Day 10]
Embryo [Day 12]
Embryo [Day 14]
Log(cell number)
12.5
10.0
7.5
5.0
2.5
B
Gene Ontology
Diapausing Embryo [Sep-Nov]
Elongated Embryo [Dec-Jan]
ribosomal large subunit biogenesis
protein N−linked glycosylation
rRNA processing
mitochondrial gene expression
mitochondrial translation
mitotic sister chromatid segregation
rRNA metabolic process
ERAD pathway
sister chromatid segregation
tRNA metabolic process
mitotic nuclear division
protein folding
response to endoplasmic reticulum stress
regulation of chromosome organization
nuclear chromosome segregation
regulation of membrane potential
synapse organization
germ cell development
cilium movement
axoneme assembly
Adj. p−value
1e−07
2e−07
3e−07
C
Gene Ontology
Embryo [Day 10]
Embryo [Day 12] + Embryo [Day 14]
tRNA aminoacylation for protein translation
nucleoside diphosphate catabolic process
rRNA metabolic process
ribonucleoprotein complex biogenesis
protein folding
mRNA processing
protein−RNA complex organization
regulation of translation
regulation of amide metabolic process
protein maturation
establishment of organelle localization
nucleobase−containing compound catabolic process
cellular component disassembly
heterocycle catabolic process
cellular nitrogen compound catabolic process
male gamete generation
cellular process involved in reproduction in multicellular organism
meiosis I cell cycle process
neuropeptide signaling pathway
ribonucleoside catabolic process
Adj. p−value
0.002
0.004
0.006
D
Capreolus capreolus
Percentage of samples (%)
S
G1
G2M
July
Sep–Nov
Dec–Jan
Bos taurus
Day 6
Day 8
Day 10
Day 12
Day 14
E
Diapausing [July]
Diapausing [Sep–Nov]
Elongated [Dec–Jan]
Pyruvate metabolism
Glutathione metabolism
Glycerophospholipid metabolism
Sphingolipid metabolism
Scaled log2(norm. counts + 1)
July
Sep–Nov
Dec–Jan

**Figure S4: Comparison of roe deer and cattle sequencing data. (A)** Principal component analysis (PCA) of roe deer and cattle embryos in relation to species, stage, ribosomal RNA (rRNA) levels and cell number. **(B-C)** Gene set enrichment analysis (GSEA) analysis of 1000 differentially expressed genes in **(B)** roe deer and **(C)** cattle upon elongation. **(D)** Proportion of samples in each embryo stage across different cell cycle phases. **(E)** Scaled counts of genes involved in “Pyruvate metabolism”, “Glutathione metabolism”, “Glycerophospholipid metabolism”, and “Sphingolipid metabolism” in roe deer embryos.

**Table S1: One-hundred and four genes upregulated upon elongation shared in both roe deer and cattle embryos.**

| | | | |
|---|---|---|---|
| A2M | FOXA1 | MDFI | SLC29A1 |
| ADAMTS1 | FTCD | MFAP2 | SLC2A10 |
| ADAMTS9 | FURIN | MFSD12 | SLC2A2 |
| AHSG | GCNT3 | MYC | SLC2A6 |
| AMIGO3 | GJB6 | NCLN | SLC35F6 |
| ANKRD13B | GMPPB | NOS2 | SLC37A2 |
| ATAD3A | GRHL3 | NUCB2 | SLC39A1 |
| BCAM | HACD2 | P4HB | SLC39A2 |
| BMP2 | HAND1 | PAQR8 | SLC44A4 |
| CA12 | HAS3 | PARVB | SLC48A1 |
| CALR | HKDC1 | PDIA4 | SLCO2A1 |
| CER1 | HMCN2 | PFKP | SPAG5 |
| CERCAM | HPDL | PKD2L1 | SURF4 |
| CHAC1 | HS3ST3B1 | PKM | TBX20 |
| CRELD2 | HSD17B1 | PLK1 | TCIRG1 |
| CRISPLD2 | ID3 | PRXL2A | TF |
| CYGB | IGF2BP1 | RBM38 | TGM1 |
| DGKZ | IRX3 | RND1 | THOP1 |
| DHCR7 | ISL2 | SATB2 | TM4SF1 |
| DKK1 | ITPKB | SCARB1 | TMBIM1 |
| DMBX1 | KAZALD1 | SEC61A1 | TMEM104 |
| DNAJB11 | KIF18B | SERPINF2 | TMEM129 |
| FADS1 | KIF22 | SLC10A1 | TMEM150B |
| FETUB | LMAN1 | SLC1A5 | VARS1 |
| FKBP10 | LPCAT3 | SLC22A23 | WNT7B |
| FKBP11 | LRRC59 | SLC27A6 | ZFP91 |
